# Three-Dimensional Network Topology of Irradiated Cancer Spheroids Revealed by Optical Coherence Tomography

**DOI:** 10.64898/2026.11.25.769092

**Authors:** Mark Arndt, Rico Hansler, Luca Tirinato, Joao Seco, Werner Nahm, Ute Schepers, Maria Francesca Spadea

**Affiliations:** Institute of Biomedical Engineering, Karlsruhe Institute of Technology (KIT), Karlsruhe, Germany; Division of Biomedical Physics in Radiation Oncology, German Cancer Research Center (DKFZ), Heidelberg, Germany; Institute of Functional Interfaces, Karlsruhe Institute of Technology (KIT), Karlsruhe, Germany; Department of Medical and Surgical Sciences, University Magna Græcia of Catanzaro, Catanzaro, Italy; Faculty of Physics and Astronomy, Ruprecht-Karls-University, Heidelberg, Germany

**Keywords:** optical coherence tomography, tumour spheroids, radiation response, graph theory, label-free imaging

## Abstract

Radiation therapy is central to cancer treatment, yet label-free quantification of tumour radiation response in three dimensions (3D) is lacking. Cell-graph topology relates to radiation response in monolayers; here we extend this to 3D tumour spheroids using optical coherence tomography. SAS spheroids were irradiated with 0–12 Gy and imaged at days 8 and 11. High-backscattering subregions from watershed segmentation served as nodes of a proximity graph, yielding eleven network metrics for *N* = 197 spheroids. Ten of eleven metrics differed between doses (Kruskal–Wallis, all *p <* 0.001): graph density rose from a median of 0.20 at 0 Gy to 0.48 at 12 Gy, clustering from 0.50 to 0.65, whereas path length fell from 2.57 to 1.59 and diameter from 5 to 3. At 0 Gy, spheroid volume (*p* = 0.021) and projected area (*p* = 0.009) were larger in 10,000-cell/well spheroids, whereas no graph metric differed (all *p* ≥ 0.21). Most dose effects, including density, persisted across broad algorithm parameter ranges; *σ* and degree assortativity were exceptions, and *σ* also depended on imaging day. Random geometric graphs with matched node counts did not reproduce the dose-dependent changes in density, clustering, path length or *σ*. Results are exploratory (single cell line and experiment).

## 1 Introduction

Radiation therapy (RT) is a primary treatment modality for a broad range of solid tumours, administered to approximately half of all cancer patients either alone or in combination with surgery and systemic therapy [1, 2]. RT exploits ionising radiation to induce lethal DNA double-strand breaks in malignant cells, ultimately causing cell death and tumour regression [2]. Despite its widespread clinical use, RT efficacy is critically hampered by *radioresistance* (RR), an innate or acquired capacity of tumour cells to withstand irradiation through mechanisms including enhanced DNA repair, autophagy, hypoxia adaptation, and cancer stem cell enrichment [3–5]. Developing quantitative, label-free methods to assess RR is therefore of considerable clinical importance for personalising RT planning and predicting treatment outcomes.

Standard assays for cellular radiosensitivity include the clonogenic colony formation assay [6, 7], neutral comet assay, and *γ*-H2AX immunofluorescence staining, which are widely used because they directly relate cell survival to radiation dose. However, these techniques are terminal, labour-intensive, and treat each colony as a monolithic entity, disregarding the intrinsic spatial heterogeneity of cells within irradiated colonies.

Beyond individual cell viability, the morphological and spatial organisation of cancer cells has recently emerged as an informative, population-level readout of radiation response. Several studies have linked radiation-induced morphological changes (cell area enlargement, altered eccentricity, and filopodia formation) to the acquisition of radioresistance in mammary and oral cancer lines [8–10]. Building on this insight, Tirinato and colleagues [11] demonstrated that the topology of two-dimensional (2D) cell networks within irradiated colonies of human lung cancer cells (H460, A549, and Calu-1) correlates with both radiation dose and intrinsic cell-line radioresistance. By positioning individual cell centres as graph nodes, connected probabilistically using the Waxman model, they showed that the small-world coefficient *σ* of the resulting cell graphs is sensitive to dose for the most radio-sensitive line (H460), while more radio-resistant lines (Calu-1) exhibit minimal topological change. These findings suggested that graph topology could serve as a quantitative indicator of cellular radiation response independently of classical morphological metrics.

A fundamental limitation acknowledged by Tirinato et al. is that their analysis was confined to 2D monolayer colonies. Planar cultures lack the diffusion gradients, cell-to-cell cooperation, and hypoxic microenvironments characteristic of solid tumours *in vivo* [12–14]. Three-dimensional tumour spheroids, by contrast, exhibit concentric proliferating, quiescent, and necrotic zones and represent a substantially more faithful *in vitro* tumour model [14]. Extending the network-topology approach to 3D spheroids is therefore a conceptually natural but technically demanding step that requires volumetric imaging of the spheroid interior.

Optical coherence tomography (OCT) offers an ideal modality for this purpose [15, 16]. OCT is a label-free, contact-free imaging technique that provides depth-resolved, micrometre-resolution images of optically scattering samples. Unlike confocal fluorescence microscopy, OCT does not require exogenous labels or thin sample preparation, and its millimetre-scale penetration depth is sufficient to image entire spheroids of typical diameter (300–600 *µ*m) in under two minutes per spheroid. OCT has previously been applied to monitor spheroid growth, necrotic core development, and drug response [17, 18], but has not been used to derive graph-theoretic topology metrics as a radiation response readout. Radiomic signatures of OCT volumes of irradiated tumour spheroids have been analysed previously [19]; the present study instead analyses cell-graph topology.

In this study, we image SAS human tongue squamous cell carcinoma spheroids with OCT after irradiation with doses ranging from 0 to 12 Gy, segment three-dimensional high-backscattering structural subregions, construct spatial proximity graphs, and systematically compute eleven network metrics. We show that the resulting 3D graph topology is sensitive to radiation dose, that graphs become denser, more clustered and shorter in path length with increasing dose (with the small-world coefficient *σ* decreasing as a secondary, parameter-sensitive metric), and that these topology changes show no association with the two different initial seeding densities used, providing no evidence that they are a seeding-density artefact.

## 2 Results

### 2.1 OCT imaging and three-dimensional graph construction

Tumour spheroids were imaged by OCT at days 8 and 11 post-irradiation across five radiation dose levels (0, 2, 4, 8, and 12 Gy) and two initial seeding densities (5,000 and 10,000 cells/well); the experimental and analysis pipeline is shown in Figure 1 (see Methods). Within each spheroid, a node was defined as a high-backscattering OCT subregion identified by watershed segmentation, and two nodes were connected by an edge when their centroid-to-centroid distance did not exceed 180 *µ*m (see Methods for all processing steps and parameters). Representative 3D graph visualisations overlaid on OCT maximum-intensity projections are shown in Figure 2.

**Figure 1:**
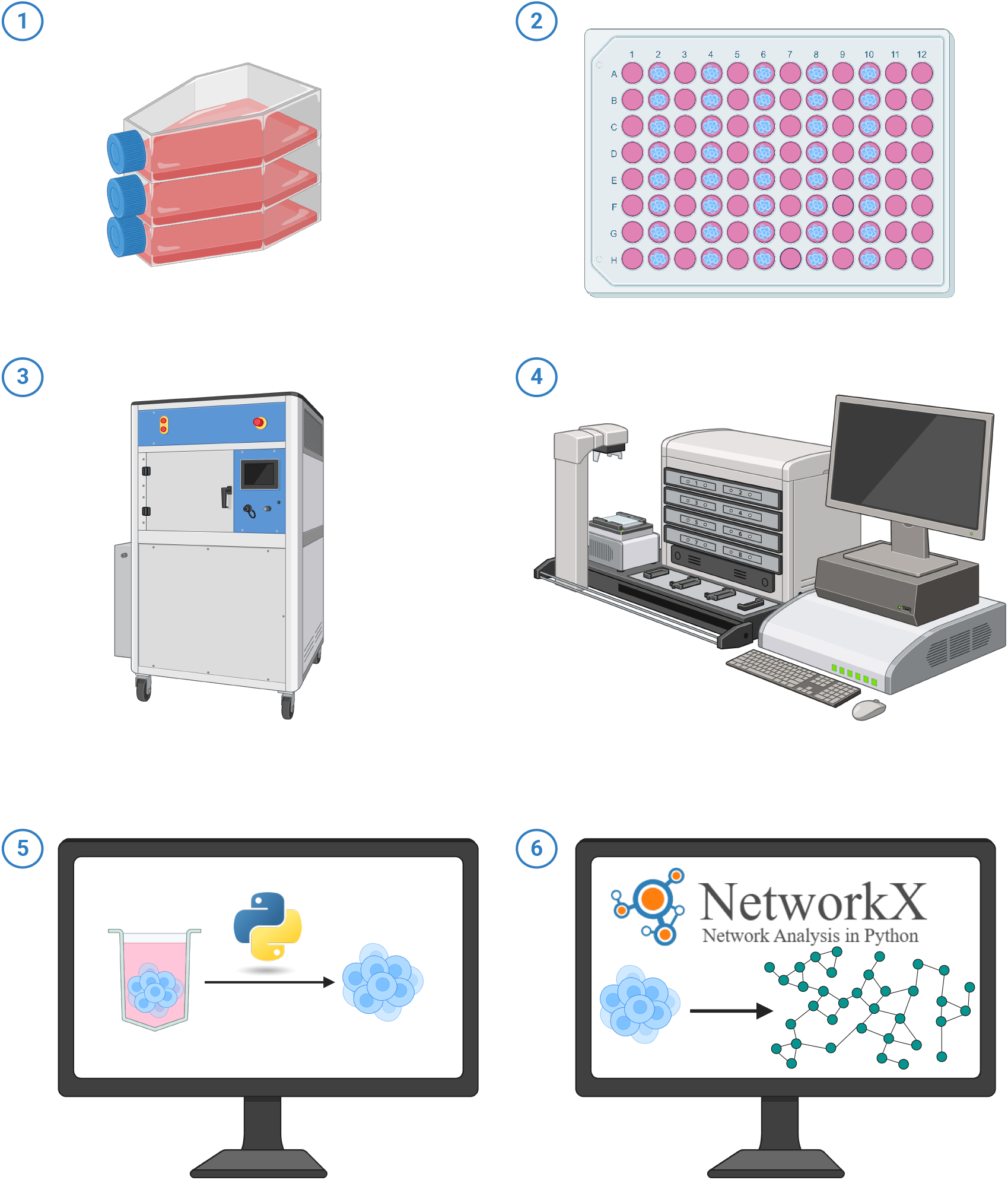
Experimental and analysis pipeline. (1) SAS tongue squamous cell carcinoma cells were expanded in culture flasks. (2) Cells were seeded into agarose-coated 96-well plates to form spheroids. (3) Spheroids were irradiated with 0, 2, 4, 8 or 12 Gy using a cabinet X-ray irradiator. (4) Spheroids were imaged by spectral-domain OCT at days 8 and 11 post-irradiation. (5) Each OCT volume was segmented in Python to obtain a spheroid mask, within which high-backscattering subregions were identified by Gaussian smoothing, peak detection and watershed segmentation. (6) Subregion centroids served as nodes of a spatial proximity graph, from which eleven network metrics were computed using NetworkX. Created with BioRender.com

**Figure 2:**
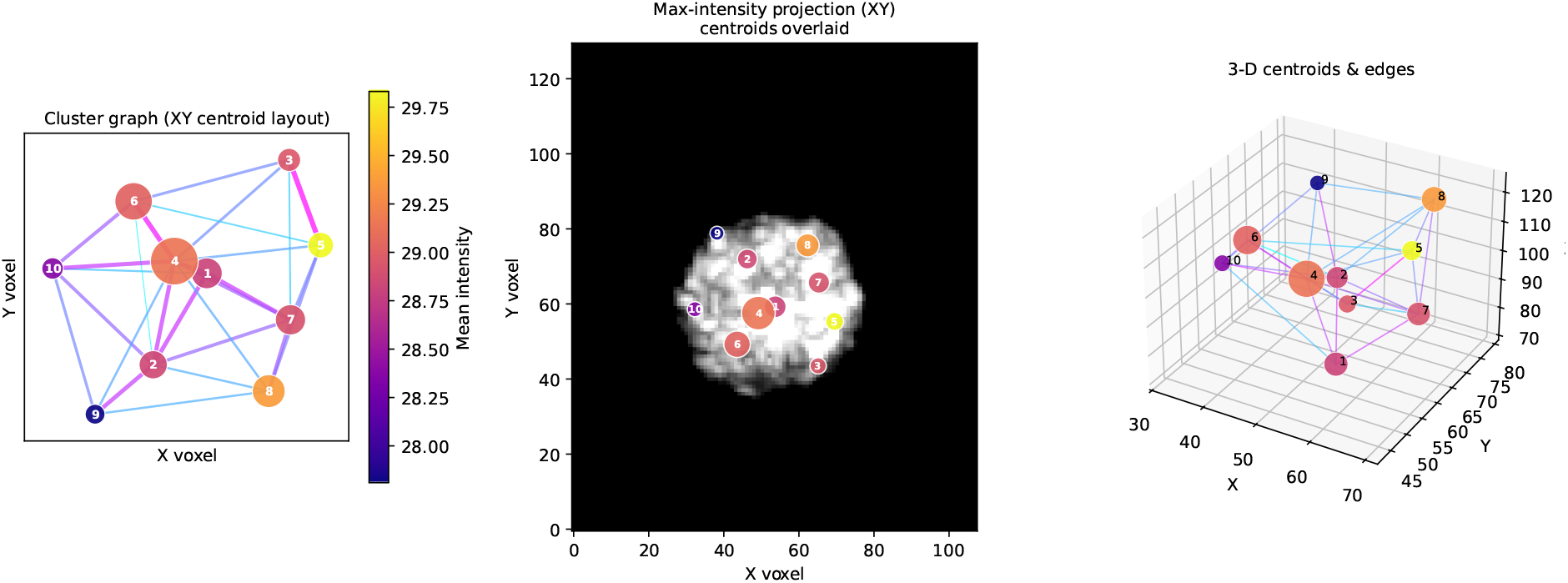
Representative 3D graph visualisation for a single spheroid. *Left*: spatial proximity graph in the XY centroid layout; node colour encodes mean OCT intensity (plasma colour map); node size scales with subregion volume; edge colour and width reflect inter-node distance. *Centre*: XY maximum-intensity projection of the OCT volume with detected centroids overlaid. *Right* : three-dimensional scatter of node centroids and edges. Shown here is a 0-Gy spheroid (5,000 cells initial seeding).

The biological identity of these subregions was not validated in this study (see Discussion). All spheroids analysed in this study were drawn from a single overall cell-culture experiment (one seeding/irradiation batch), imaged across five independently prepared replicate 96-well plates at day 8 and four at day 11. In total, 290 wells had a usable intensity/mask pair across all doses and both seeding densities. Of these, 91 (31%) were excluded because the resulting graph was disconnected (more than one connected component) or a specular reflection artefact was visible in the volume and a further 2 were excluded as node-count outliers by the 1.5×IQR criterion, leaving *N* = 197 spheroids for the dose-response analysis (day 8: 93; day 11: 104; 0 Gy: *n* = 35; 2 Gy: *n* = 30; 4 Gy: *n* = 31; 8 Gy: *n* = 37; 12 Gy: *n* = 64).

Across these spheroids, the number of detected nodes (high-backscattering subregions) ranged from a few to several dozen depending on dose and spheroid size, and the resulting graphs exhibited a broad range of connectivity patterns (Figure 2). Eleven network metrics were computed per spheroid: number of nodes and edges, graph density, average degree, average local clustering coefficient, transitivity, average path length, diameter, average betweenness centrality, degree assortativity, and the small-world coefficient *σ* (see Methods for definitions).

### 2.2 Radiation dose systematically modulates 3D network topology

A Kruskal–Wallis (KW) test across the five dose groups revealed that 10 of 11 network metrics differed significantly with radiation dose (all *p <* 0.001; Table 1; Figure 3). These included: number of nodes, graph density, average degree, average clustering coefficient, transitivity, average path length, diameter, average betweenness centrality, degree assortativity, and the small-world coefficient *σ*. Only the number of edges was not modulated significantly (*p* = 0.318), consistent with the graph becoming both smaller (fewer nodes) and denser with dose in a way that leaves the raw edge count comparatively stable while density (edges normalised by the number of possible node pairs) increases sharply.

**Figure 3:**
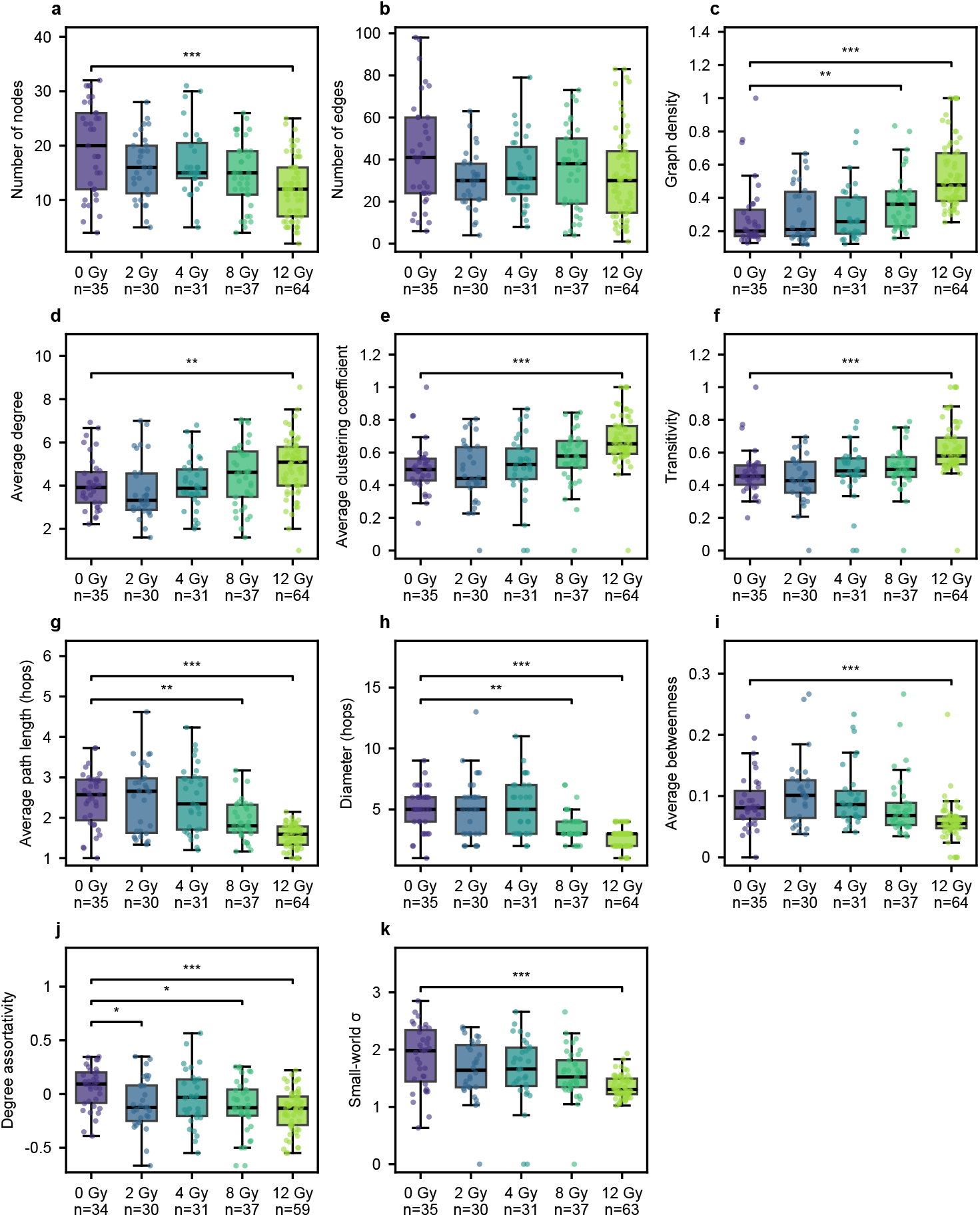
Graph metrics across radiation dose groups. Panels a–k show the eleven graph metrics (a: number of nodes; b: number of edges; c: density; d: average degree; e: average clustering coefficient; f: transitivity; g: average path length; h: diameter; i: average betweenness centrality; j: degree assortativity; k: small-world coefficient *σ*) for the 0, 2, 4, 8 and 12 Gy groups; the number of spheroids per group is given in the *x*-axis tick labels (*N* = 197 in total); the *y*-axes of average path length and diameter are in hops (edges), and the other metrics are dimensionless (counts of nodes and edges, or ratios). Boxes span the interquartile range (IQR) with the line at the median; whiskers extend to the most extreme values within 1.5×IQR of the box; points are individual spheroids. Brackets show post-hoc two-sided Mann–Whitney U comparisons of each dose vs. 0 Gy with Bonferroni-adjusted *p*-values (4 comparisons; * *p <* 0.05, ** *p <* 0.01, *** *p <* 0.001; ns, *p* ≥ 0.05). Kruskal–Wallis *p*-values for all metrics are given in Table 1. Both seeding densities and days 8 and 11 post-irradiation are pooled. The small-world coefficient *σ* (panel k) decreases from a median of 1.979 at 0 Gy to 1.304 at 12 Gy; individual spheroids in each group can fall below 1.

**Table 1:** Graph metrics by radiation dose group, given as median [Q1–Q3] (Q = quartile), with Kruskal–Wallis *p*-values (column *p*) across the five dose groups (*α* = 0.05). *n*: number of spheroids per group; both seeding densities and days 8 and 11 are pooled. Spheroids with an undefined value of a metric were omitted for that metric only: degree assortativity *n* = 34, 30, 31, 37 and 59 (0, 2, 4, 8 and 12 Gy) and *σ n* = 63 at 12 Gy; all other rows use the group sizes shown. Post-hoc comparisons of each dose vs. 0 Gy used Mann–Whitney U tests with Bonferroni correction (see Figure 3).

| Metric | 0 Gy<br>(*n*=35) | 2 Gy<br>(*n*=30) | 4 Gy<br>(*n*=31) | 8 Gy<br>(*n*=37) | 12 Gy<br>(*n*=64) | *p* |
|---|---|---|---|---|---|---|
| Nodes | 20<br>[12–26] | 16<br>[11.25–20] | 15<br>[14–20.5] | 15<br>[11–19] | 12<br>[7–16] | < 0.001 |
| Edges | 41<br>[24–60] | 30<br>[21–38] | 31<br>[23.5–46] | 38<br>[19–50] | 30<br>[14.75–44] | 0.318 |
| Density | 0.200<br>[0.170–0.329] | 0.210<br>[0.170–0.436] | 0.257<br>[0.184–0.403] | 0.362<br>[0.228–0.440] | 0.477<br>[0.382–0.670] | < 0.001 |
| Avg. degree | 3.92<br>[3.21–4.63] | 3.32<br>[2.88–4.56] | 3.88<br>[3.48–4.75] | 4.62<br>[3.48–5.58] | 5.08<br>[4.00–5.80] | < 0.001 |
| Avg. clustering | 0.496<br>[0.429–0.563] | 0.441<br>[0.388–0.632] | 0.527<br>[0.437–0.625] | 0.578<br>[0.507–0.671] | 0.653<br>[0.592–0.762] | < 0.001 |
| Transitivity | 0.455<br>[0.403–0.521] | 0.427<br>[0.354–0.543] | 0.488<br>[0.457–0.564] | 0.497<br>[0.450–0.571] | 0.579<br>[0.528–0.690] | < 0.001 |
| Avg. path length | 2.57<br>[1.94–2.94] | 2.65<br>[1.63–2.97] | 2.34<br>[1.71–3.00] | 1.80<br>[1.63–2.32] | 1.59<br>[1.33–1.78] | < 0.001 |
| Diameter | 5<br>[4–6] | 5<br>[3–6] | 5<br>[3–7] | 3<br>[3–4] | 3<br>[2–3] | < 0.001 |
| Avg. betweenness | 0.081<br>[0.063–0.108] | 0.101<br>[0.064–0.126] | 0.086<br>[0.066–0.108] | 0.068<br>[0.053–0.089] | 0.055<br>[0.047–0.067] | < 0.001 |
| Assortativity | 0.094<br>[−0.082, 0.202] | −0.124<br>[−0.250, 0.080] | −0.030<br>[−0.204, 0.136] | −0.127<br>[−0.202, 0.042] | −0.132<br>[−0.288, −0.023] | < 0.001 |
| *σ* | 1.979<br>[1.442–2.340] | 1.640<br>[1.341–2.080] | 1.662<br>[1.364–2.034] | 1.522<br>[1.346–1.814] | 1.304<br>[1.222–1.495] | < 0.001 |

Several consistent trends with increasing dose were observed (Table 1). First, graph density more than doubled (0.20 to 0.48) and the average clustering coefficient increased (0.50 to 0.65) while average path length decreased substantially (2.57 to 1.59) and diameter decreased from 5 to 3; together, these changes indicate progressive network compaction: the detected subregions become more tightly interconnected (Figure 3). Second, the small-world coefficient *σ* decreased from a median of 1.98 at 0 Gy to 1.30 at 12 Gy, a reduction of 34%, but this metric was significantly different from control only at 12 Gy, showed a significant dose × day interaction (*p* = 0.009), and was among the metrics most sensitive to the graph-construction parameters (see the robustness analysis below); it is therefore regarded as a less robust, secondary metric. Average degree increased in parallel (3.9 to 5.1), and average betweenness centrality decreased (0.081 to 0.055), consistent with the shorter path lengths in smaller, denser graphs. Third, degree assortativity declined with dose and changed sign (0.094 at 0 Gy to −0.13 at 12 Gy), reflecting a shift from a network in which well-connected nodes preferentially link to other well-connected nodes toward a mildly disassortative, more random connectivity pattern at high doses. The number of detected nodes also decreased with dose (median nodes: 20 at 0 Gy to 12 at 12 Gy). Post-hoc comparisons against the 0 Gy control (Mann–Whitney U, Bonferroniadjusted for four comparisons) showed that most of this dose-response was concentrated at the higher doses: density (adjusted *p* = 0.0023), average path length (*p* = 0.0016), diameter (*p* = 0.0010), and degree assortativity (*p* = 0.014) already differed from control at 8 Gy, whereas number of nodes, average degree, average clustering, transitivity, average betweenness, and *σ* differed only at 12 Gy (all adjusted *p* ≤ 0.0085; *σ*: *p <* 0.001). Degree assortativity additionally differed from control at 2 Gy (*p* = 0.048) but not at 4 Gy (*p* = 0.38), and the number of edges did not differ from control at any dose (all adjusted *p* ≥ 0.29). The dose-response was therefore gradual rather than a step change (Figure 3).

To assess whether this dose effect was consistent across the experimental design rather than driven by a subset of it, the KW test was repeated separately within each measurement day, within each seeding-density group, and within each of the nine individual replicate plates (in this paragraph, “significant” denotes KW *p <* 0.05 within the stratum). The dose effect remained significant in both day 8 and day 11 considered separately for all metrics except number of edges (and, within day 8 alone, average degree), and largely so in both the 5,000- and 10,000-cells/well seeding-density groups considered separately (with three metric-level exceptions at the *p <* 0.05 threshold across these strata: average degree was not significant within day 8 alone, *p* = 0.096; degree assortativity was not significant within the 10,000-cells/well group alone, *p* = 0.093; and number of edges, not significant overall or within the 10,000-cells/well group, was nominally significant within the 5,000-cells/well group alone, *p* = 0.029). These are the expected consequence of the roughly two-to five-fold smaller per-group sample sizes available within each stratum relative to the pooled analysis, rather than evidence of a qualitatively different dose response in any subgroup: the corresponding pooled or other-stratum test was significant (respectively, non-significant, for number of edges) in every case. Within individual replicate plates (*n* = 10–29 spheroids per plate, split across five dose groups), the dose effect was still detectable for most metrics in the larger plates, but several metrics lost significance in the smallest plates (*n* = 10–17); this pattern is consistent with reduced statistical power from the correspondingly small per-dose sample sizes within a plate (as few as 0–2 spheroids per dose group in the smallest plates; the day-8 plate 3 contained no 2-Gy spheroid) rather than a plate-specific reversal of the effect. A two-way, rank-based Scheirer–Ray–Hare test for dose × day, dose × seeding-density, and dose × plate interaction effects found no significant interaction for any metric except a significant dose × day interaction for *σ* (*p* = 0.009) and a significant dose × seeding-density interaction for number of edges (*p* = 0.043, the one metric with no significant overall dose effect); the dose × plate interaction was non-significant for all eleven metrics (*p* = 0.12–0.90), with no evidence that the dose effect depends on which replicate plate a spheroid was drawn from, although some plate × dose cells were sparse or empty. All other interaction tests were non-significant (*p* = 0.051–0.91). Together, these analyses provide no evidence that the reported dose effects are an artefact of pooling across days, seeding densities, or replicate plates, with the caveat that the day-dependence of the *σ* dose-response should be interpreted with some caution.

### 2.3 Graph topology shows no association with seeding-density-driven size differences

A potential alternative explanation for the observed topological changes is that radiation simply shrinks spheroids, and smaller networks inherently exhibit different metric values regardless of true structural reorganisation. To test this, we compared graph metrics as well as spheroid volume and maximum XY projected area between the two seeding-density groups (5,000 and 10,000 cells/well) at 0 Gy, where any differences would arise purely from the initial seeding density rather than from radiation effects (Table 2; Figures 4 and 5 ).

**Figure 4:**
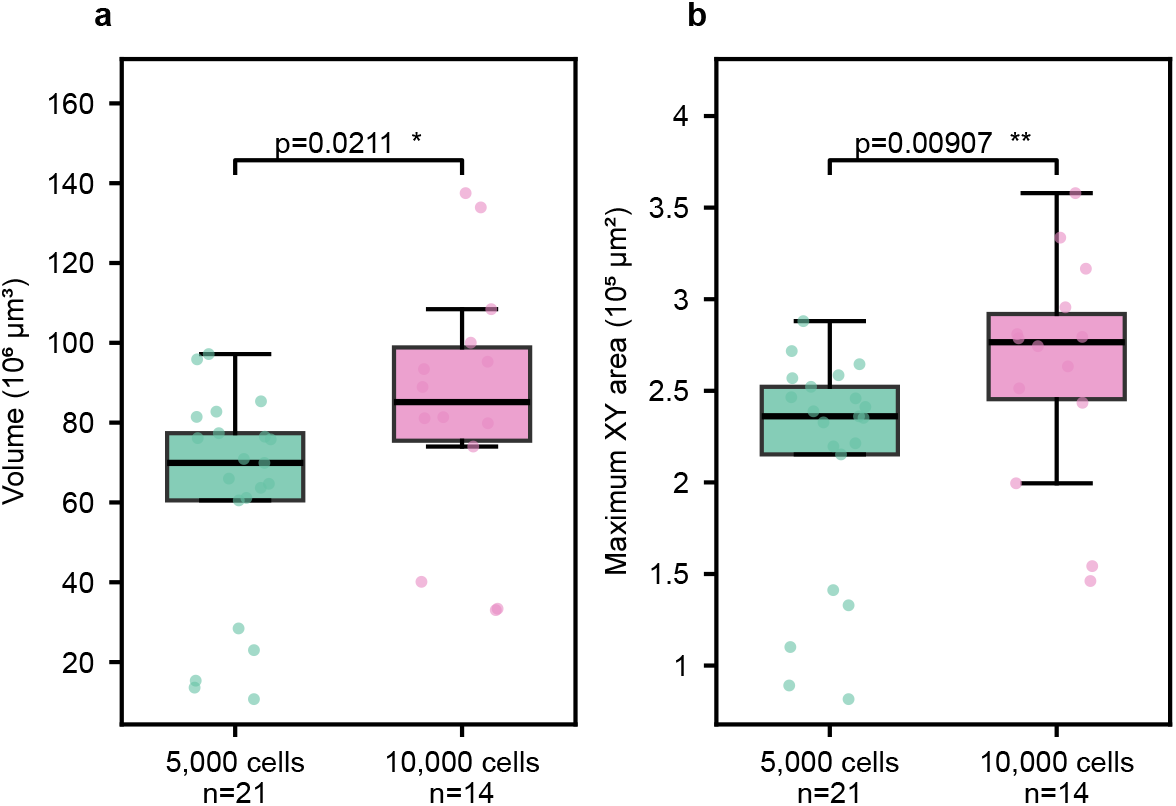
Spheroid size at 0 Gy: 5,000 vs 10,000 cells/well. Box plots (median, IQR box, whiskers at 1.5×IQR, points individual spheroids) of (a) spheroid volume (10^6^ *µ*m^3^) and (b) maximum XY projected area (10^5^ *µ*m^2^) between spheroids formed at 5,000 cells/well (*n* = 21) and 10,000 cells/well (*n* = 14; *N* = 35 total), restricted to the unirradiated (0-Gy) condition. Brackets indicate two-sided Mann–Whitney U test results (ns, *p* ≥ 0.05). Both spheroid volume (*p* = 0.021) and maximum XY area (*p* = 0.009) differed significantly between the two seeding-density groups in this matched-dose comparison, with the 10,000-cell/well group larger in both measures, as expected from the higher seeding density. All eleven graph topology metrics were nonetheless not significantly different between groups (all *p* ≥ 0.21; Figure 5).

**Figure 5:**
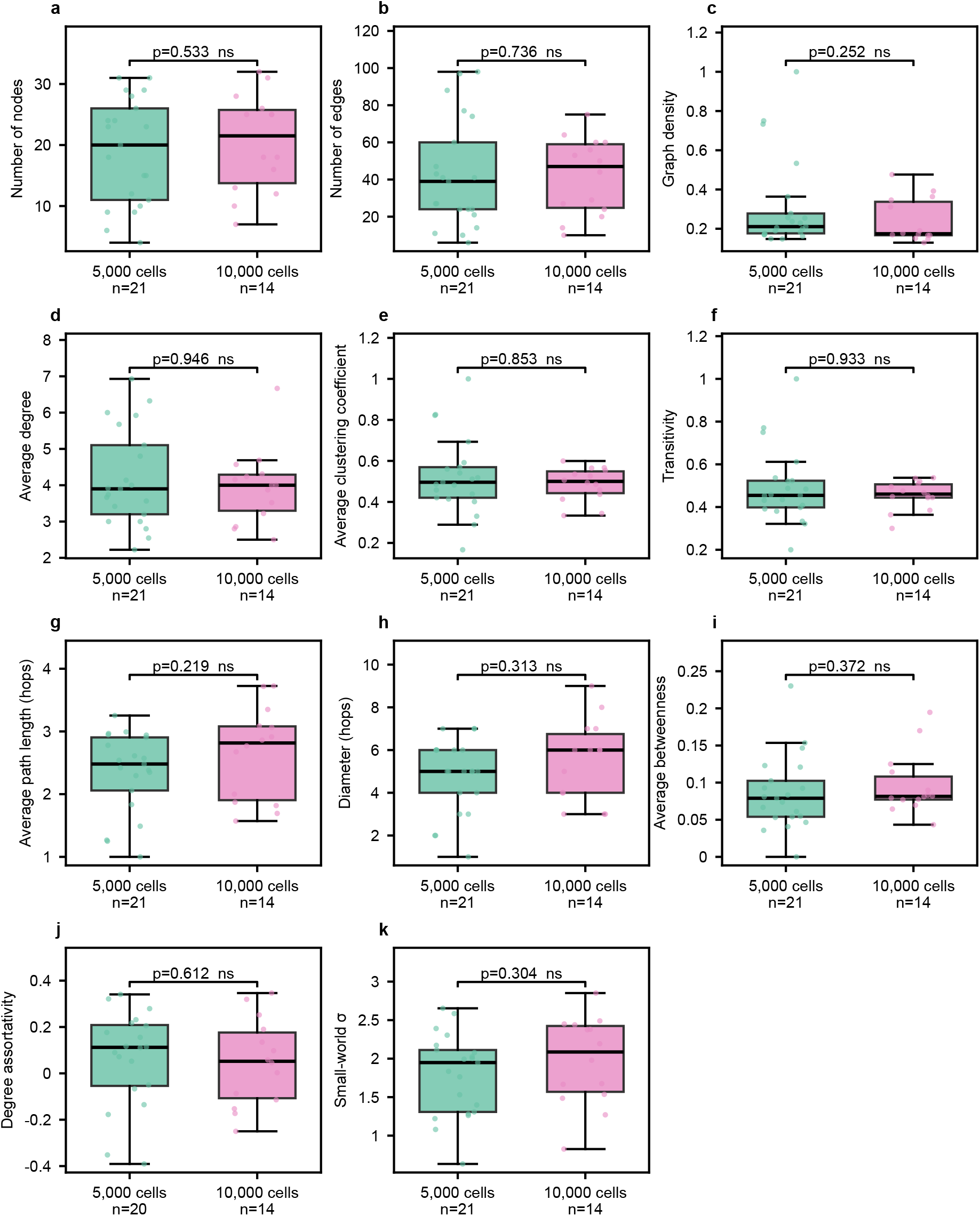
Graph topology at 0 Gy: 5,000 vs 10,000 cells/well seeding-density groups. Panels a–k show the eleven graph metrics (same order as in Figure 3) for spheroids formed at 5,000 cells/well (*n* = 21) and 10,000 cells/well (*n* = 14; *N* = 35 total), restricted to the unirradiated (0-Gy) condition. Boxes show the median and IQR, whiskers extend to 1.5×IQR, and points are individual spheroids. Exact *p*-values of two-sided Mann–Whitney U tests are printed in each panel (ns, *p* ≥ 0.05); none of the eleven metrics differed significantly between groups (all *p* ≥ 0.21), despite spheroid volume and projected area both being significantly larger in the 10,000-cell/well group at this same dose (see Figure 4), indicating that graph topology does not simply track spheroid size.

**Table 2:**
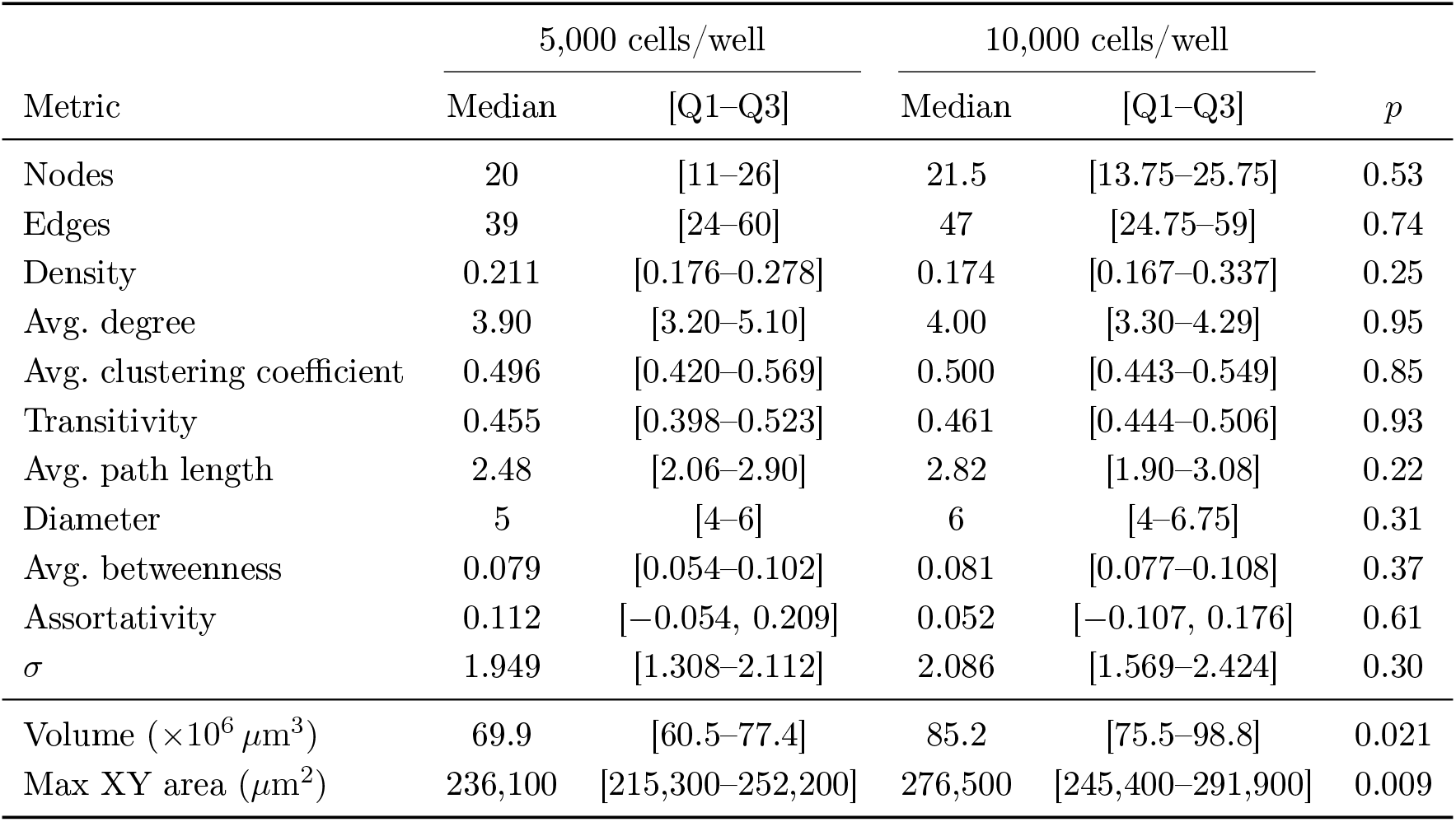
Graph metrics and morphology at 0 Gy: 5,000-cell/well vs 10,000-cell/well seeding-density groups. Values are median [Q1–Q3] (Q = quartile); *n*_5k_ = 21, *n*_10k_ = 14 (*N* = 35); for degree assortativity *n*_5k_ = 20 (one undefined value omitted; *n*_10k_ = 14). *p*: exact *p*-value of the two-sided Mann–Whitney U test between the two groups.

|  | 5,000 cells/well |  | 10,000 cells/well |  |  |
|---|---|---|---|---|---|
| Metric | Median | [Q1–Q3] | Median | [Q1–Q3] | *p* |
| Nodes | 20 | [11–26] | 21.5 | [13.75–25.75] | 0.53 |
| Edges | 39 | [24–60] | 47 | [24.75–59] | 0.74 |
| Density | 0.211 | [0.176–0.278] | 0.174 | [0.167–0.337] | 0.25 |
| Avg. degree | 3.90 | [3.20–5.10] | 4.00 | [3.30–4.29] | 0.95 |
| Avg. clustering coefficient | 0.496 | [0.420–0.569] | 0.500 | [0.443–0.549] | 0.85 |
| Transitivity | 0.455 | [0.398–0.523] | 0.461 | [0.444–0.506] | 0.93 |
| Avg. path length | 2.48 | [2.06–2.90] | 2.82 | [1.90–3.08] | 0.22 |
| Diameter | 5 | [4–6] | 6 | [4–6.75] | 0.31 |
| Avg. betweenness | 0.079 | [0.054–0.102] | 0.081 | [0.077–0.108] | 0.37 |
| Assortativity | 0.112 | [−0.054, 0.209] | 0.052 | [−0.107, 0.176] | 0.61 |
| *σ* | 1.949 | [1.308–2.112] | 2.086 | [1.569–2.424] | 0.30 |
| Volume (×10<sup>6</sup> *μ*m<sup>3</sup>) | 69.9 | [60.5–77.4] | 85.2 | [75.5–98.8] | 0.021 |
| Max XY area (*μ*m<sup>2</sup>) | 236,100 | [215,300–252,200] | 276,500 | [245,400–291,900] | 0.009 |

In this matched-dose comparison (*n*_5k_ = 21, *n*_10k_ = 14), spheroid volume (*p* = 0.021) and maximum XY projected area (*p* = 0.009) both differed significantly between the two seeding densities (Mann–Whitney U test, two-sided; Figure 4), with medians of 69.9 × 10^6^ vs 85.2 × 10^6^ *µ*m^3^ for volume and 236,100 vs 276,500 *µ*m^2^ for projected area, both larger in the 10,000-cell/well group, as expected from the higher initial seeding density. This confirms that the two seeding-density groups differ in spheroid size even at matched (0 Gy) dose. None of the eleven graph metrics differed significantly between the two seeding-density groups at 0 Gy (all *p* ≥ 0.21; Figure 5; Table 2), including the small-world coefficient *σ* (*p* = 0.30, medians: 1.95 vs 2.09), graph density (*p* = 0.25), average clustering coefficient (*p* = 0.85), and average path length (*p* = 0.22). Because spheroid size did differ significantly between the two seeding densities in this comparison, the absence of any corresponding difference in graph topology strengthens, rather than merely being consistent with, the interpretation that the dose-dependent topological changes reported above reflect genuine radiation-induced reorganisation of the spheroid interior rather than a simple consequence of spheroid size.

### 2.4 Dose effects on graph topology are not explained by node number

Radiation reduced the number of detected subregions (median 20 nodes at 0 Gy versus 12 at 12 Gy), so some metrics could change simply because smaller graphs have different topology. To test this, we generated random geometric graphs that kept the real node count of every spheroid but no other structure: nodes were placed uniformly at random in a common volume, under the same minimum separation and edge rule as the real graphs (Methods). We repeated this 100 times and tested the five dose groups of each repetition exactly as for the real data. By construction, the random graphs reproduce the dose dependence of node number (*p* = 2.1 × 10*^−^*^5^) and, as a consequence, of edge number (median null KW *p* = 2.5 × 10*^−^*^5^), average degree (6.1 × 10*^−^*^5^) and average betweenness (6.0 × 10*^−^*^4^). In contrast, density, clustering, transitivity, path length, diameter and *σ* did not differ between dose groups in most repetitions (median null KW *p* = 0.11, 0.18, 0.31, 0.46, 0.43 and 0.18; *p <* 0.05 in 30, 22, 12, 10, 10 and 24% of repetitions), whereas the real data gave *p* = 6.1×10*^−^*^12^, 1.3×10*^−^*^9^, 3.7×10*^−^*^10^, 9.7×10*^−^*^12^, 4.9 × 10*^−^*^12^ and 7.2 × 10*^−^*^7^. The same held for 12 Gy versus 0 Gy: density *p* = 1.7 × 10*^−^*^9^ (real) versus 0.084 (null median), clustering 5.4 × 10*^−^*^9^ versus 0.16, path length 1.8 × 10*^−^*^9^ versus 0.57, and *σ* 1.2 × 10*^−^*^6^ versus 0.14.

The edge count showed the opposite pattern. Node number alone predicts fewer edges at high dose (null *p* = 2.5 × 10*^−^*^5^), yet the real edge count did not differ between dose groups (*p* = 0.32). High-dose spheroids therefore have more edges per node than node number predicts, which is the denser connectivity seen in the other metrics. Degree assortativity was intermediate (real *p* = 7.6 × 10*^−^*^4^, null median 0.061).

### 2.5 Graph topology findings are robust to algorithm parameter choices

To assess whether the observed dose-dependent topology changes depend critically on the specific algorithm parameters, we performed a one-at-a-time (OAT) sensitivity analysis in which the relative intensity threshold (0.60–0.80), the minimum peak distance (30–72 *µ*m), and the edge distance threshold (90–300 *µ*m) were each varied independently while the other two were held at their baseline values (Figure 6). In this section, “significant” denotes a KW *p <* 0.05 for the dose effect under the parameter set considered (the *p*-values are shown in Figure 6).

**Figure 6:**
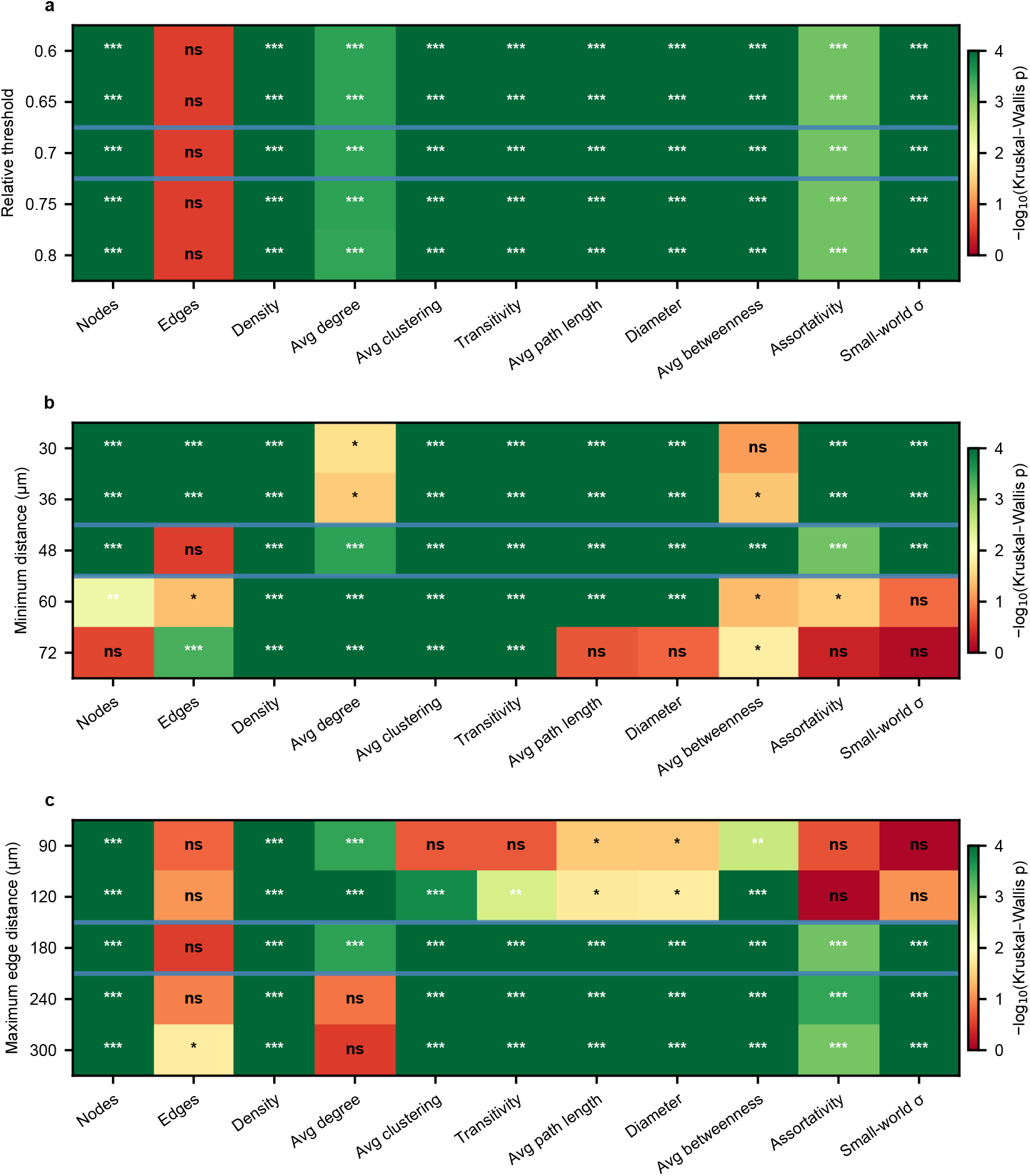
Parameter robustness of dose-group significance. Kruskal–Wallis (KW) *p*-values for all eleven graph metrics under one-at-a-time variation of three algorithm parameters: (a) relative intensity threshold, (b) minimum peak distance (*µ*m), and (c) edge distance threshold *d*_max_ (*µ*m) (y-axis labels in the panels: “Relative threshold” = relative intensity threshold, “Minimum distance (*µ*m)” = minimum peak distance, and “Maximum edge distance (*µ*m)” = edge distance threshold *d*_max_), computed on the *N* = 197 analysis set with the physically isotropic pipeline (Methods). Colour encodes − log_10_ of the KW *p*-value on a red–yellow–green scale (green = smaller *p*, dark red ≈ *p* = 1), capped at − log_10_ *p* = 4; cells print stars for the significance level (∗ *p <* 0.05, ** *p <* 0.01, *** *p <* 0.001) and “ns” (not significant) for *p* ≥ 0.05. The baseline value of each parameter is the row framed by the two blue lines. Number of edges is the only metric non-significant at baseline throughout the relative intensity threshold sweep and for most of the maximum-edge-distance sweep, consistent with it being the one metric not significantly dose-modulated overall (Table 1). The minimum peak distance sweep shows the largest parameter sensitivity, with several metrics losing significance only at the two most extreme (sparsest-graph) values tested; average degree additionally loses significance at the two largest edge distance threshold values.

The significance pattern was completely insensitive to the intensity threshold: all ten metrics that were significant at baseline (all except number of edges, which remained non-significant throughout) retained exactly the same significance pattern across the full tested range (relative intensity threshold 0.60–0.80; Figure 6a). This is because the relative intensity threshold applies a relative floor to the peak amplitude; within this range the same local maxima are detected regardless of the specific value.

The minimum peak distance was the most influential parameter, consistent with it directly controlling how many watershed seeds are placed: median node count per spheroid fell from 62 (0 Gy) / 28 (12 Gy) at a minimum peak distance of 30 *µ*m to 6 / 5 at a minimum peak distance of 72 *µ*m (Figure 6b). This was also the only parameter for which the identity of the non-significant metric changed: at the two smallest values (30, 36 *µ*m) average degree was only nominally significant while number of edges was significant; at the baseline (48 *µ*m) this reverses (average degree strongly significant, number of edges not), matching the pattern reported for the rest of the study (Table 1), whereas at 60 and 72 *µ*m number of edges was again significant (nominally at 60 *µ*m). Graph density, average clustering, and transitivity remained significant across the entire tested range (30–72 *µ*m). At the two largest values (60, 72 *µ*m), where the median node count per spheroid falls to ≈8–12 and 5–6 respectively, several path- and distance-based metrics lost significance: at a minimum peak distance of 72 *µ*m, number of nodes, average path length, diameter, degree assortativity, and *σ* were no longer significant. At 60 *µ*m, *σ* was already non-significant, and at 30 *µ*m average betweenness was non-significant. The baseline value of 48 *µ*m lies within the stable region for ten of eleven metrics (all but number of edges, which is non-significant at baseline); only the two most extreme increases (60, 72 *µ*m) degrade the topological metrics that depend on having enough nodes to resolve path- and distance-based structure. The edge distance threshold (*d*_max_) showed a graded rather than sharply thresholded pattern (Figure 6c). At the two smallest values (90, 120 *µ*m), the graphs are comparatively sparse and the more structure-dependent metrics lose significance first: at *d*_max_ = 90 *µ*m, average clustering, transitivity, degree assortativity, and *σ* are not significant, and average path length and diameter are only nominally significant; at *d*_max_ = 120 *µ*m, average clustering and transitivity recover significance but degree assortativity and *σ* remain non-significant. From *d*_max_ = 180 *µ*m (baseline) through 300 *µ*m, number of nodes, density, average clustering, transitivity, average path length, diameter, average betweenness, degree assortativity, and *σ* are significant throughout; average degree is the exception, significant at the baseline value (180 *µ*m) but not at the two larger values tested (240 and 300 *µ*m). This pattern is opposite to its behaviour under the minimum peak distance, and we have no mechanistic explanation for it beyond the graph becoming close to fully connected at these larger distance thresholds. Graph density remains significant across the entire tested range because it normalises edge count by *n*(*n* − 1) and so continues to track relative connectivity differences even in very sparse or near-complete graphs. Overall, the baseline of *d*_max_ = 180 *µ*m lies at the boundary of, rather than comfortably within, the informative range for the most topologically complex metrics (degree assortativity and *σ* in particular only become and remain significant from this value upward), and results for these two metrics should be considered somewhat more sensitive to this parameter than the other metrics.

Overall, this analysis confirms that the core dose-response finding is not an artefact of a single arbitrary parameter choice: the majority of metrics remain significant across most of the tested parameter space, and where sensitivity does exist (the small-world coefficient and degree assortativity under a smaller edge distance threshold, average degree under a larger edge distance threshold, or several metrics under a much larger minimum peak distance) it is concentrated at the more extreme ends of each tested range rather than at the baseline parameters used throughout this study.

## 3 Discussion

In this work, we show that the 3D network topology of tumour spheroids, derived from OCT structural volumes and analysed using graph-theoretic metrics, is systematically altered by ionising radiation. Ten of eleven computed network metrics differed significantly across doses of 0–12 Gy (all KW *p <* 0.001), with networks becoming denser, more clustered and shorter in path length (density 0.20 to 0.48, clustering 0.50 to 0.65, path length 2.57 to 1.59) and the small-world coefficient *σ* decreasing as a secondary, parameter-sensitive metric (1.98 to 1.30 at 12 Gy), and this dose effect was largely consistent when tested separately within each measurement day and each seeding-density group. A seeding-density comparison at matched (0 Gy) dose confirmed that spheroid volume and projected area were both significantly larger in the 10,000-cell/well group (*p* = 0.021 and *p* = 0.009), yet none of the eleven graph metrics differed significantly between the two seeding densities (all *p* ≥ 0.21), indicating that the reported dose effects on graph topology are not simply a consequence of spheroid size. Random geometric graphs with the same node counts also failed to reproduce the dose-dependent changes in density, clustering, path length, diameter and *σ*, indicating that these changes are not explained by the reduction in node number alone. Together, these findings support OCT-derived 3D network topology as a dose-dependent structural readout of irradiated tumour spheroids in this single-cell-line, single-experiment, exploratory setting.

The present results can be compared directly with the 2D cell-graph approach. The current study extends the cell-graph topology framework introduced by Tirinato et al. [11] from 2D monolayer colonies to 3D spheroids. Both studies report dose-dependent changes in graph topology, and both include the small-world coefficient among the metrics examined. However, the direction of the *σ* change differs: Tirinato et al. observed an *increase* in *σ* with dose for the radio-sensitive H460 line (reaching *σ* ≈ 1.8 at 8 Gy), whereas we report a *decrease* in *σ* with dose (from 1.98 to 1.30 at 12 Gy). This apparent discrepancy may reflect a fundamental difference in the definition of graph nodes. In Tirinato et al., nodes are individual cells; the Waxman-connected cell graph captures cell-to-cell spatial relationships at single-cell resolution, and the emergence of *σ >* 1 at high doses signals the formation of small clusters of surviving cells in a depleted colony. In the present study, nodes are large OCT-derived structural subregions each encompassing many cells; the proximity graph captures the meso-scale architectural organisation of the spheroid interior. With increasing dose, fewer subregions were detected and the remaining nodes formed denser, shorter networks with higher clustering. Higher clustering and shorter path lengths accompany this compaction, whereas *σ*, which is normalised to a random reference graph, decreases with dose; since random graphs with the same node counts did not reproduce this decrease (median null *p* = 0.18), it cannot be attributed to the lower node number alone. At every dose level tested, the median *σ* remained *>* 1, although individual spheroids occasionally fell below 1.

The biological meaning of the graph nodes requires careful qualification. The graph nodes analysed here are OCT intensity-peak subregions identified by watershed segmentation of backscatter contrast, not independently identified cells, cell populations, or histologically defined structural units. In OCT, signal magnitude at a given voxel is proportional to local optical backscattering, which is governed by spatial variations in refractive index at the scale of the illumination wavelength [16]. One plausible, but in the present study unvalidated, interpretation is that regions of elevated backscatter within a tumour spheroid correspond to locally denser cellular assemblies or organelle-rich cytoplasm, both of which could produce stronger refractive-index contrast with surrounding interstitial fluid [20]. Under this interpretation, the graph nodes would represent structurally dense micro-environments within the spheroid, and the proximity edges their spatial organisation, with radiation-induced cell death and tissue disruption altering the number and spatial distribution of these micro-environments and hence the graph. The increase in graph density and clustering coefficient with dose coincided with a decrease in the number of detected subregions. This pattern is compatible with, but does not establish, a more compact arrangement of high-backscattering regions in irradiated spheroids. Establishing what the segmented subregions actually correspond to at the cellular level would require correlating OCT-derived graphs with an independent radiobiological readout (e.g. viability or proliferation staining) on matched or serial sections, which was not performed here and is an important direction for future validation. Future work using speckle-variance (SV) OCT, which provides dynamics-sensitive contrast reflecting intracellular and intercellular motion [21], could likewise provide a complementary, biologically richer readout that helps separate structural compaction from metabolic activity.

A further motivation for the approach is its three-dimensional and non-destructive nature. Extending the topology analysis to three dimensions is biologically important because the spatial distribution of viable, quiescent, and necrotic cells in a spheroid is inherently 3D [12]. A 2D projection or a single focal plane would systematically misrepresent the true 3D connectivity of the structural network. OCT is particularly well suited to this task because it is non-destructive, label-free, and fast (1–2 min per spheroid), making it compatible with longitudinal designs and with high-throughput screening workflows.

Several limitations of the current study should be noted. First, the analysis was performed on a single cell line (SAS); validating the approach across cell lines with known differential radiosensitivity, as performed in the 2D setting by Tirinato et al., is a necessary next step to demonstrate clinical utility. Second, the graph construction relies on parameters (Gaussian smoothing *σ_G_*, minimum peak distance, edge distance threshold) that were held constant across all samples. A one-at-a-time sensitivity analysis varying each parameter across a broad range confirmed that most significant metrics remain significant across reasonable parameter choices, with sensitivity concentrated at the extremes of each tested range (Section 2.5; Figure 6). Third, the geometric proximity edge model used here differs from the probabilistic Waxman model of Tirinato et al.; direct numerical comparisons of absolute metric values across the two frameworks should be made with caution, although the qualitative topological trends are directly comparable. Fourth, all spheroids analysed here were drawn from a single overall cell-culture experiment: the dose-group comparisons pool multiple replicate plates and both measurement days, and while the dose effect was largely consistent when tested separately within each day, within each seeding-density group, and within each replicate plate, and no significant dose × day, dose × seeding-density, or dose × plate interaction was detected for any metric except *σ* (dose × day, *p* = 0.009) and number of edges (dose × seeding-density, *p* = 0.043; the one metric with no overall dose effect) (Section 2.2), independent biological replication (separate seeding/irradiation experiments performed on different occasions) was not performed and remains an important next step before these findings can be considered generalisable. Fifth, the segmented structural subregions are identified purely from OCT backscatter contrast and have not been independently validated against a cellular- or viability-level ground truth (see above); the biological identity of these subregions therefore remains an open question.

## 4 Methods

An overview of the experimental and analysis pipeline is provided in Figure 1.

### 4.1 Cell culture

SAS human tongue squamous cell carcinoma cells were provided by Dr. Ina Kurth (German Cancer Research Center, DKFZ, Heidelberg, Germany). Cells were maintained in Dulbecco’s Modified Eagle Medium (DMEM) supplemented with 10% fetal bovine serum (FBS) and 1% penicillin/streptomycin at 37 °C in a humidified 5% CO_2_ atmosphere. Passages 17–19 were used for all experiments.

### 4.2 Spheroid formation

Spheroids were formed by seeding 5,000 or 10,000 cells per well in 100 µl of culture medium in 96-well agarose plates. Agarose plates were prepared with 50 µl of 1.5% agarose solution per well. After seeding, spheroids were maintained for a further six days to reach the target size prior to irradiation.

### 4.3 Irradiation

Spheroids were irradiated using a cabinet-based photon (X-ray) irradiator (Faxitron MultiRad 225 [22]) at doses of 0, 2, 4, 8, or 12 Gy, at a dose rate of approximately 2 Gy min*^−^*^1^ at the sample position. OCT imaging was performed at days 8 and 11 post-irradiation. A cross-sectional design was adopted: at each measurement day, several independently prepared 96-well replicate plates (identical seeding and culture protocol, from a single overall cell-culture experiment) were imaged and then not reused for the other day. All dose groups were irradiated on the same day and imaged at identical post-irradiation time points (days 8 and 11); every dose group contributes spheroids to both days.

Each plate followed the same layout, with 5,000-cell/well spheroids and 10,000-cell/well spheroids in separate row sets (four rows each) and the five dose conditions (0, 2, 4, 8, 12 Gy) assigned to dedicated columns within each row set, giving up to 40 wells (5 doses × 2 seeding densities × 4 rows) per plate. Day 8 was imaged across 5 such replicate plates and day 11 across 4 replicate plates (see Results for the resulting per-dose sample sizes after quality filtering).

### 4.4 Optical coherence tomography

OCT is a label-free, non-contact imaging technique that provides depth-resolved cross-sectional and volumetric images of biological samples [15, 16]. Coherence gating of backscattered light from a broadband source enables micrometre-scale axial ranging without the need for physical sectioning or exogenous labels. Raster scanning in two lateral directions yields a complete 3D intensity volume.

Spheroid imaging was performed using a spectral-domain OCT system (Thorlabs Ganymede GAN612C1/M [23]), fitted with an OCT-LK4-BB objective, with a centre wavelength of 880 nm operating at a 5 kHz A-scan rate. This platform and objective combination has been independently characterised in-house at an axial resolution of 4.5 *µ*m (in tissue) and a lateral resolution of 12 *µ*m. Spheroids were maintained at roughly 37 °C on a temperature-controlled stage during acquisition. Imaging time per spheroid was approximately 1–2 min. Each acquisition covered a field of view of 0.9 mm × 0.9 mm (lateral) × 3.63 mm (axial), large enough to contain the full spheroid footprint plus a surrounding rim of clear medium and the entire spheroid depth. Volumetric data were exported and analysed as linear backscattered intensity: no logarithmic (dB) compression, depth-dependent sensitivity roll-off correction, or explicit attenuation compensation was applied prior to segmentation or graph construction. Resulting volumetric data were stored as NIfTI files with voxel dimensions of 6.0 × 6.0 × 3.5 *µ*m^3^ (XY × Z). **Mask generation and quality control.** Binary segmentation masks delineating the spheroid boundary were generated per volume using an interactive, semi-automated procedure. The mean intensity volume (mean over repeated frames where applicable) was Otsu-thresholded [24] within a manually defined axial range, followed by binary morphological closing and opening (spherical structuring element, 1-voxel radius) and removal of connected components smaller than 3,000 voxels; the largest remaining connected component was retained as the spheroid mask. Residual out-of-plane medium or debris was removed via interactively adjustable axial and lateral crop boundaries. Each mask was visually inspected against the underlying intensity volume, and only masks judged to faithfully delineate the spheroid boundary were accepted into the analysed dataset; volumes with ambiguous or failed segmentation were excluded prior to graph construction (see also the connectivity- and outlier-based exclusion criteria below). Accepted masks were stored as co-registered NIfTI files.

### 4.5 Image segmentation and graph construction

All processing was implemented in Python 3 using NumPy [25], SciPy [26], scikit-image [27], and NetworkX [28]. Because the OCT voxel grid is anisotropic (6.0 × 6.0 × 3.5 *µ*m^3^), every step below that involves a spatial distance or separation criterion was carried out in physical (*µ*m) units rather than raw voxel-index units, as detailed for each step.

#### Smoothing and peak detection

Within each binary spheroid mask, the OCT intensity volume was smoothed with a 3D Gaussian filter with a physically isotropic standard deviation of 9 *µ*m in every dimension (*σ_G_* = 1.5 voxels in XY, *σ_G_* = 2.57 voxels in Z, i.e. scaled by the voxel-size ratio 6.0*/*3.5). Smoothed voxels outside the mask were set to zero. Local intensity maxima were detected with peak_local_max from scikit-image on a version of the smoothed volume resampled to an isotropic voxel grid (Z resampled to match the XY pitch), enforcing a minimum peak distance of 8 voxels (48 *µ*m), measured in true 3D physical distance, and a relative intensity threshold of 70% of the in-mask maximum; detected peaks were then mapped back onto the original (anisotropic) voxel grid for watershed seeding.

#### Watershed segmentation

Detected peaks served as seeds for a 3D watershed transform applied to the negated smoothed volume within the spheroid mask, yielding a label image in which each voxel was assigned to the nearest intensity peak region. The centroid of each labelled region constituted a graph node; node attributes included region size (number of voxels) and mean OCT intensity. Subregion sizes substantially exceeded the system’s speckle correlation length, making speckle noise an unlikely explanation for the detected structures.

#### Graph construction

An undirected graph was constructed by connecting any two nodes whose Euclidean centroid-to-centroid distance, computed in physical (*µ*m) coordinates from the anisotropic voxel geometry, did not exceed an edge distance threshold of *d*_max_ = 30 XY-voxel-equivalents (≈180 *µ*m). Edge weights were set to this physical distance. This geometric proximity model differs from the probabilistic Waxman model used by Tirinato et al. [11]; the threshold was chosen to ensure that topologically meaningful connections are formed across the typical inter-node distances observed in the spheroids while avoiding spurious long-range connections.

Spheroids for which the resulting graph contained more than one connected component were excluded from analysis. Additional exclusions applied the 1.5×IQR outlier criterion to the node-count distribution; excluded spheroids had either far fewer (fragmented segmentation) or far more (over-segmentation artefacts) nodes than the bulk of the dataset.

### 4.6 Graph metric computation

Eleven network metrics were computed per spheroid using NetworkX.

1. Number of nodes *n* and edges *m*.
2. **Graph density**: *ρ* = 2*m /* [*n*(*n* − 1)].
3. **Average degree**: ⟨*k*⟩ = 2*m/n*.
4. **Average local clustering coefficient** ⟨*C*⟩: mean over all nodes *i* of *C_i_* = 2*E_i_ /* [*k_i_*(*k_i_* − 1)], where *k_i_* is the degree of node *i* and *E_i_* the number of existing edges among its *k_i_* neighbours.
5. **Transitivity** (global clustering coefficient): *T* = 3 × (closed triangles)*/*(connected triples).
6. **Average path length** ⟨*ℓ*⟩: mean shortest path length over all pairs of nodes in the largest connected component, computed using NetworkX’s average_shortest_path_length.
7. **Diameter** *d*: maximum shortest path length.
8. **Average betweenness centrality** ⟨*b*⟩: fraction of shortest paths passing through each node, averaged.
9. **Degree assortativity** *r*: Pearson correlation of degrees at both ends of each edge.
10. Small-world coefficient *σ* [29]:

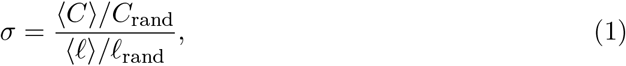

where the Erdős–Rényi random-graph references are *C*_rand_ ≈ ⟨*k*⟩*/*(*n* − 1) and *ℓ*_rand_ ≈ ln *n/* ln⟨*k*⟩ [30]. Networks with *σ >* 1 are classified as small-world.

Two of these metrics can be undefined for individual spheroids and were then recorded as missing and omitted from the analysis of that metric only. Degree assortativity was recorded as missing whenever NetworkX raised an error or returned no value, which occurs when it is undefined, e.g. when all nodes have the same degree (zero degree variance). *σ* was recorded as missing when the random-graph path-length reference *ℓ*_rand_ was undefined (average degree ⟨*k*⟩ ≤ 1) or when ⟨*ℓ*⟩ could not be computed.

### 4.7 Parameter sensitivity analysis

A one-at-a-time (OAT) sensitivity analysis was conducted to assess robustness of the dose-group comparisons to algorithm parameter choices. Three parameters were varied independently while the remaining two were held at their baseline values: the relative intensity threshold ∈ {0.60, 0.65, 0.70, 0.75, 0.80}, the minimum peak distance ∈ {30, 36, 48, 60, 72} *µ*m, and the edge distance threshold *d*_max_ ∈ {90, 120, 180, 240, 300} *µ*m. For each of the 13 parameter sets, all eleven network metrics were recomputed for every spheroid in the statistical analysis set (days 8 and 11; *N* = 197 spheroids) and the Kruskal–Wallis test was repeated. To minimise computation, the 3D Gaussian-smoothed volume was computed once per spheroid and reused across all threshold and distance sweeps; for the *d*_max_ sweep, the watershed node centroids from the baseline run were additionally reused and only the graph was rebuilt.

### 4.8 Statistical analysis

All statistical analyses were performed in Python with SciPy [26]. Dose-response analyses were performed on spheroids imaged at days 8 and 11 post-irradiation, at which radiation effects on spheroid architecture were expected to have developed. All spheroids analysed in this study were derived from a single overall cell-culture experiment (one seeding/irradiation batch), with each measurement day represented by several independently prepared replicate 96-well plates (see Irradiation, above); the unit of analysis throughout is one OCT-imaged well (spheroid). The total numbers of replicate plates and analysed wells are reported in the Results. Because all data derive from a single experiment, the reported effects should be interpreted as an exploratory, single-experiment demonstration pending independent biological replication.

#### Choice of tests

The graph metrics are bounded or discrete (e.g. density, clustering coefficients, diameter, node and edge counts) or skewed, and the dose groups were of unequal size (30–64 spheroids per group). Normality was assessed with the Shapiro–Wilk test for each metric within each dose group; normality was rejected at *α* = 0.05 (uncorrected) in 31 of the 55 metric × dose-group combinations. Rank-based tests were therefore used instead of analysis of variance.

The Kruskal–Wallis (KW) test was used to assess whether each graph metric differed across the five dose groups, pooling both seeding densities and both time points as the primary analysis. Post-hoc comparisons between each irradiated dose group and the 0-Gy control used the two-sided Mann–Whitney U (MW) test with Bonferroni correction for four comparisons. To assess whether the pooled dose effect was consistent across the experimental design, the KW test was repeated separately within each measurement day (day 8 only; day 11 only), within each seeding-density group (5,000 and 10,000 cells/well), and within each of the nine individual replicate plates, and a two-way, rank-based Scheirer–Ray–Hare test [31] was used to test for dose × day, dose × seeding-density, and dose × plate interaction effects on each metric (plate identity defined per measurement day, since plate numbers were not unique across days). The comparison between the 5,000-cell/well and 10,000-cell/well seeding-density groups was performed with the two-sided MW test on the 0-Gy dose group. Spheroid volume and projected XY area were compared between seeding-density groups at 0 Gy only, using the same two-sided MW test. The significance level was *α* = 0.05 (all Mann–Whitney tests were two-sided); exact *p*-values are reported in the text and tables (values below 0.001 are given as *p <* 0.001), and asterisks in figures denote *, *p <* 0.05; **, *p <* 0.01; ***, *p <* 0.001 (Bonferroni-adjusted for the post-hoc comparisons); ns (not significant) denotes *p* ≥ 0.05. Box plots show the median (line), the interquartile range (box), whiskers extending to the most extreme observation within 1.5×IQR of the box, and individual spheroids as overlaid points.

### 4.9 Random geometric null model

For every spheroid of the final analysis set, one random geometric graph was generated with the same number of nodes as the real graph, so that each dose group retained its real node-count distribution (medians 20, 16, 15, 15 and 12 at 0, 2, 4, 8 and 12 Gy). Nodes were placed uniformly at random inside a sphere of median volume, by random sequential addition with a minimum pairwise separation of 48 *µ*m, the separation used by the peak detector. Edges were created between nodes closer than 180 *µ*m, and the same eleven metrics were computed as for the real graphs. Disconnected graphs were redrawn, as disconnected spheroids were excluded from the real analysis. This procedure was repeated 100 times.In each repetition the five dose groups were compared with the Kruskal–Wallis test, and each dose with 0 Gy using two-sided Mann–Whitney U tests with Bonferroni correction, as for the real data. We report the median *p*-value over the repetitions and the fraction of repetitions with *p <* 0.05. Because node counts were fixed to the real values, the null reproduces the node-count comparison exactly.

### 4.10 Use of AI tools

Large language models were used to polish and refine the language of the manuscript and to assist with coding basic functions. All content, analyses and results were reviewed and verified by the authors, who take full responsibility for the work.

## Data Availability

The processed graph metric data supporting the findings of this study are available from the corresponding author upon reasonable request. OCT imaging data are available upon request subject to data-sharing agreements.

## Acknowledgements

The authors thank Dr. Ina Kurth (DKFZ, Heidelberg) for providing the SAS cell line and for the use of the Faxitron MultiRad 225 irradiator.

## Author contributions

M.A. conceived the study, performed OCT imaging, conducted the graph analysis, and wrote the manuscript. R.H. cultivated the cells. W.N. reviewed the manuscript and contributed to discussions on OCT. L.T. reviewed the manuscript and contributed to discussions on the methodology. U.S. provided the laboratory space for cell culture and supervised the work. J.S. supervised the work (radiobiology). M.F.S. supervised the work, contributed to discussions on the methodology, and reviewed the manuscript.

